## Supplementary Materials for "Structural insights into scaffold-guided assembly of the Pseudomonas phage D3 capsid"

Anna K. Belford, *et al.*

**This PDF file includes:**

Supplementary Text
Figs. S1 to S9
Tables S1 to S5
Movies S1 to S4
References (55 to 72)

**Other Supplementary Materials for this manuscript include the following:**

Movies S1 to S4

### Supplementary Text

### Supplementary Methods – Short Descriptions (SMS#)

Supplementary Method 1.

***Buffers Related to this Study***

| | |
|---|---|
| *Lysis buffer*: | 50 mM Tris-HCl pH 8.0, 5 mM EDTA |
| *D3 P1 Lysis buffer*: | 50 mM Tris-HCl pH 6.8, 100 mM K Glutamate, 10 mM EDTA |
| *G buffer*: | 20 mM Tris-HCl pH 7.5, 100 mM NaCl |
| *D3 P1 buffer*: | 50 mM Tris/50 mM Bis-Tris Propane-HCl pH 6.0, 100 mM potassium glutamate |
| *Dialysis buffer*: | 5 mM Tris-HCl pH 7.5 |
| *D3 P1 Dialysis buffer*: | 2.5 mM Tris/2.5 mM Bis-Tris Propane pH 6.0 |
| *TAMg buffer*: | 40 mM Tris-HCl, 20 mM acetic acid pH 8.1, 1 mM $MgSO_4$ |

Supplementary Method 2.

***Creation of expression plasmids and strains containing phage D3 capsid genes.*** Phage D3 was obtained from Andrew Kropinski. pT7-5 (*55*) was used as the expression plasmid for D3 capsid genes. Plasmids used for expression contained the T7 promoter, D3 capsid genes 4, 5, and/or 6, and an ampicillin resistance gene. Plasmids were created by cut-and-paste digestion, purification, and ligation; site directed mutagenesis in which 2 of 4 primers are phosphorylated to allow for ligation and reamplification (*56*); Klenow DNA polymerase I fill-in (Boehringer Mannheim, Ingelheim, Germany); and PCR amplification with pfu (and later phusion) PCR kits (New England Biolabs, Ipswich, MA) (ST2 & SMD1). Created plasmids were co-transformed with pLysS into BL21(DE3) by electroporation (SMD2).

Supplementary Method 3.

***Production of Proheads from Plasmids.***

Proheads were purified through PEG precipitation, differential sedimentation, velocity centrifugation with glycerol gradients, and anion exchange chromatography. BL21(DE3) pLysS strains containing expression plasmids with phage D3 capsid genes (ST3) were grown and expressed under autoinduction conditions (*57*). Cells were harvested by centrifugation, resuspended in lysis buffer, and lysed by adding 0.25% (w/v) Triton X-100. Viscosity was reduced by treating with 25 μg/mL deoxyribonuclease I and 12.5 mM $MgSO_4$. The lysate was clarified by centrifugation and precipitated with 6% (w/v) PEG 8000/0.5 M Salt (potassium glutamate for P1; NaCl for P2). Precipitate was pelleted by centrifugation, resuspended in buffer (D3 P1 buffer for P1; G for P2), and repelleted by ultracentrifugation in Ti-45 (Beckman Coulter, Brea, CA) at 35000 rpm for 2 hours. Pellet was resuspended, loaded onto 10-30% glycerol gradients in same buffer, and sedimented in SW32Ti (Beckman) for 105 minutes at 29000 rpm. The prohead band was extracted, dialyzed, ran over an anion exchange column (Applied Biosystems, Waltham, MA), and pelleted again in Ti-45 for 2 hours at 35000 rpm. Pellet was resuspended in a small amount of buffer.

Supplementary Method 4.

***Small Scale Expression of Capsid Genes from Plasmids***

Expression of plasmids under autoinduction conditions (*57*) allows for rapid evaluation of plasmid expression phenotypes of capsid genes from (ST2) when performed small scale as compared to large-scale preps (1.5 mL vs. 300 mL) (*58*). After growth to high density, cells were lysed with Triton X-100, treated with DNase I, and separated by centrifugation into cell pellet and cell lysate. The cell lysate was separated into two fractions, one of which was PEG precipitated, pelleted, and resuspended. The three fractions of cell pellet (PT), cell lysate (SUP), and PEG pellet (PEG) were analyzed by native agarose and SDS polyacrylamide gel electrophoresis.

Supplementary Method 5.

***Gel Electrophoresis.***

Native Agarose Gel Electrophoresis

Agarose gel electrophoresis was performed in submarine gel boxes (Hoeffer, Holliston, MA) using TAMg buffer. Samples were loaded with a dye/glycerol loading solution (50% (v/v) glycerol, 0.025% (w/v) bromophenol blue, 0.025% (w/v) xylene cyanol XFF) and run at ~120 V on agarose gels (usually 1.2% (w/v)). Gels were stained with ~0.002% (w/v) Coomassie Brilliant Blue R250 in ~4.5% (v/v) methanol, ~9.9% acetic acid, then destained with 10% acetic acid, and photographed wet.

Samples were TCA (trichloroacetic acid) precipitated prior to sodium dodecyl sulfate (SDS) polyacrylamide gel electrophoresis to prevent spontaneous crosslinking of D3 major capsid protein during sample preparation. Precipitates were washed with acetone, suspended in SDS sample buffer, heated in a boiling bath for 2.5 min, and run on 12% SDS acrylamide minigels. The original buffers described by Laemmli (*59*) were used. A low-crosslinker recipe (33.5% (w/v) acrylamide + 0.3% (w/v) methylene bisacrylamide) (*60*) was used. Gels were stained with 0.5% (w/v) Coomassie Brilliant Blue R250 in 50% (v/v) Methanol and 10% acetic acid, destained with 10% acetic acid, and photographed wet.

### Supplementary Methods – Detailed Descriptions (SMD#)

Supplementary Method 1.

***Inserting Phage D3 Capsid Proteins into pT7-5 Plasmid***

Phage D3 was obtained from Andrew Kropinski. D3 capsid proteins were PCR amplified from phage genome in 50 µL volume reactions using pfu (and later phusion) protocol (New England Biolabs, Ipswich, MA) with the primers listed in (ST1) and methods in (ST2). Vectors and inserts were digested with restriction enzymes (New England Biolabs) and ligated with T4 ligase (New England Biolabs). Site directed mutagenesis was achieved by phosphorylating 2 of 4 primers to allow ligation and amplification (*56*). Ligated plasmids were transformed into electrocompetent DH10B cells.

Supplementary Method 2.

***Transformation of Plasmids into Electro-competent Cells***

0.66 μL of single plasmid in DH10B cells, or 0.3 μL of plasmid and 0.3 μL of pLysS in BL21(DE3), were added together on ice, flicked 5-6 times, and added into electro cuvette. Cuvettes were shocked and then grown in SOC (New England Biolabs) for 45-60 min at 37°C in incubator shaking at 250rpm. Culture was plated onto appropriate selection plates and colonies grown overnight at 37°C. For plasmid purification, several colonies were picked, cultured in LB ampicillin overnight, and DNA harvested (Qiagen, Hilden, Germany). The plasmid DNA was then sent for Sanger sequencing (Genewiz, Southplainfield, New Jersey) using sequencing primers (ST1).

Supplementary Method 3.

***Production of Proheads from Plasmids***

Co-transformed strains of BL21(DE3) (ST3) were grown in 5 mL LB $amp^{50}/cam^{25}$ overnight. The culture was diluted 1:300 into TYM-5052 $amp^{50}/cam^{25}$ in a Fernbach flask and grown for 24 hours at 37°C shaking at 300 rpm in a shaker with 1-inch throw (*57*). Cells were chilled for 30 min on ice. Samples were kept cold from this point forward unless explicitly stated. Cells were collected by centrifugation in JA-16.25 (Beckman Coulter, Brea, CA) at 6k rpm for 10 min, and resuspended on ice in 60 mL Lysis buffer for Prohead 2 and D3 P1 Lysis buffer for Prohead 1. Resuspended cells were transferred to 250 mL flask, put on ice, and 1.2 mL 10% (w/v) Triton X-100 was added dropwise. While stirring, cells were cycled by heating to 23-24°C, holding for 5 min at temperature and chilling for 5 min on ice. 40 mL more of lysis buffer was added and cycled again. An additional 0.8 mL 10% Triton was added and cycled again. 1 mL of 10% Triton was added and cycled while heating to 27-28°C before holding. A further 50 mL of lysis buffer was added along with 1 mL 10% Triton. By this point, the cells would have lysed and be very viscous. After lysis, 2 mL of 1M $MgSO_4$ was added dropwise and 2 mL of 1 mg/mL DNase I was added. Lysed cells were heated to 28°C and held for 3 min or until viscosity dropped then chilled on ice for 5 min. Cell lysate was clarified by centrifugation in JA-16.25 at 8k rpm for 12 min, and PEG precipitated with 7% (w/v) PEG 8000 and 0.5 M salt (NaCl for P2, potassium glutamate for P1), and put on ice for 20 min. Precipitant was pelleted in JA-16.25 at 8k rpm for 16 min and resuspended in G buffer for P2 and D3 P1 buffer for P1. Resuspended pellet was further centrifuged in JA-16.25 at 8k rpm for 12 min to further remove any remaining insoluble material. Afterwards, the supernatant was centrifuged in Ti-45 (Beckman) at 35k rpm for 2 hr, and resultant pellet was covered with appropriate buffer (enough to cover pellet while tube stands upright). Pellet was allowed to resuspend slowly overnight. Resuspended pellet was clarified further to remove insoluble material in JA-18 (Beckman) at 8 k rpm for 10 min. This supernatant was loaded onto 10-30% glycerol gradients made with Gradient Master (Biocomp Instruments, New Brunswick, CA) in same buffer, and sedimented in SW-32Ti (Beckman) at 29k rpm for 105 min. The prohead band was extracted and dialyzed in dialysis buffer for P2 and D3 P1 dialysis buffer for P1 at room temperature. The dialyzed material was run over a POROS HQ20 anion exchange column using BIOCAD 700E perfusion chromatography system (Applied Biosystems, Waltham, MA). Collected peaks were pelleted in Ti-45 at 35k rpm for 2 hr and covered in appropriate buffer (enough buffer to cover the pellet while centrifuge tube is tilted on its side to maximize concentration). The pellet was allowed to resuspend slowly overnight.

Supplementary Method 4.

***Small Scale Expression of Capsid Genes from Plasmids***

Co-transformed strains of BL21(DE3) were grown in 2 mL minimal media MDG $amp^{50}/cam^{25}$ overnight at 37°C shaking at 250 rpm with 2 inch throw (*57*). 1.5 mL of TYM-5052 $amp^{50}/cam^{25}$ in a 18x150 mm test tube was inoculated with 5 µL of MDG overnight and incubated at 37°C shaking at 250 rpm for 24 hr (*57*). The cells were poured into 1.5 mL eppendorf tubes and harvested by centrifugation in a micro-centrifuge at 7krpm for 2 min (cold, if possible). The cells were resuspended in 300 µL of Lysis buffer for P2 or D3 P1 Lysis buffer for P1 by vortexing violently until fully resuspended. 4.5 µL of 10% Triton X-100 was added and the cells vortexed briefly and put on dry ice for 10 min. The cells were thawed slowly at room temperature for 15 min. 12 µL of 1M $MgSO_4$ and 9 µL of 1 mg/mL deoxyribonuclease I (in 1mM HCl) were added and mixed into the cells then heated at 30°C for 3 min or until viscosity dropped. Cell lysate was clarified by centrifugation at15k rpm for 10 min (cold, if possible). The insoluble fraction was resuspended with 300 µL G or D3 P1 buffer and saved for analysis. 50 µL of cell lysate was set aside for analysis. 250 µL of cell lysate was PEG precipitated with 250 µL 12% (w/v) PEG 8000/1M NaCl for P2 or 66 µL 3M K Glutamate and 79µL 30% (w/v) PEG 8000 for P1, and put on ice for 20 min. Precipitate was pelleted in micro-centrifuge at 15k rpm for 10 min in cold, and resuspended in 250 µL G or D3 P1 buffer for P2 and P1, respectively.

Supplementary Method 5.

***Gel Electrophoresis***

*Native Agarose Gel Electrophoresis*

0.36 g of LE agarose was melted in 30 mL of TAMg [40 mM Tris-HCl, 20 mM acetic acid pH 8.1, 1 mM $MgSO_4$] and poured into a mold. The gel was put into a submarine gel box (Hoeffer, Holliston, MA). 4 µL of sample was mixed 1 µL dye [50% (v/v) glycerol, 0.025% (w/v) bromophenol blue, 0.025% (w/v) xylene cyanol XFF] and 5 µL of TAMg. 4 µL was loaded onto the gel, and ran at ~120 V for 1 hr. The gels were stained with ~0.002% (w/v) Coomassie Brilliant Blue R250 in ~4.5% (v/v) methanol, ~9.9% acetic acid overnight, destained with 10% acetic acid, and photographed wet.

*Sodium Dodecyl Sulfate Polyacrylamide Gel Electrophoresis*

Samples were TCA (trichloroacetic acid) precipitated (*61*) by adding 25 µL 100% (w/v) TCA + 20 mg/mL deoxycholate (*62*) into diluted sample (10 µL sample + 225 µL 5 mM Tris-HCl pH 7.5), vortexing, and placing on ice for 10 min. Precipitate was pelleted in micro-centrifuge at max speed for 10 min. Pellet was suspended with 0.5 mL 90% (v/v), 10 mM Tris-HCl pH 7.5 at room temperature, and pelleted in micro-centrifuge at max speed. The supernatant was aspirated off, and the pellet dried under vacuum for 5 min. Pellet was suspended with 40 µL SDS sample buffer (62.5 mM Tris-HCl pJ 6.8, 2% (w/v) SDS, 10% (v/v) glycerol, 5% (v/v) Beta-mercaptoethanol) by vortexing, and heated in a boiling bath for 2.5 min. SDS samples were loaded and run on 12% SDS polyacrylamide minigel using original buffers described by Laemmli (*59*). A low-crosslinker recipe (33.5% (w/v) acrylamide + 0.3% (w/v) methylene bisacrylamide) (*60*) was used. Gels were stained with 0.5% (w/v) Coomassie Brilliant Blue R250 in 50% (v/v) Methanol and 10% acetic acid, destained with 10% acetic acid, and photographed wet.

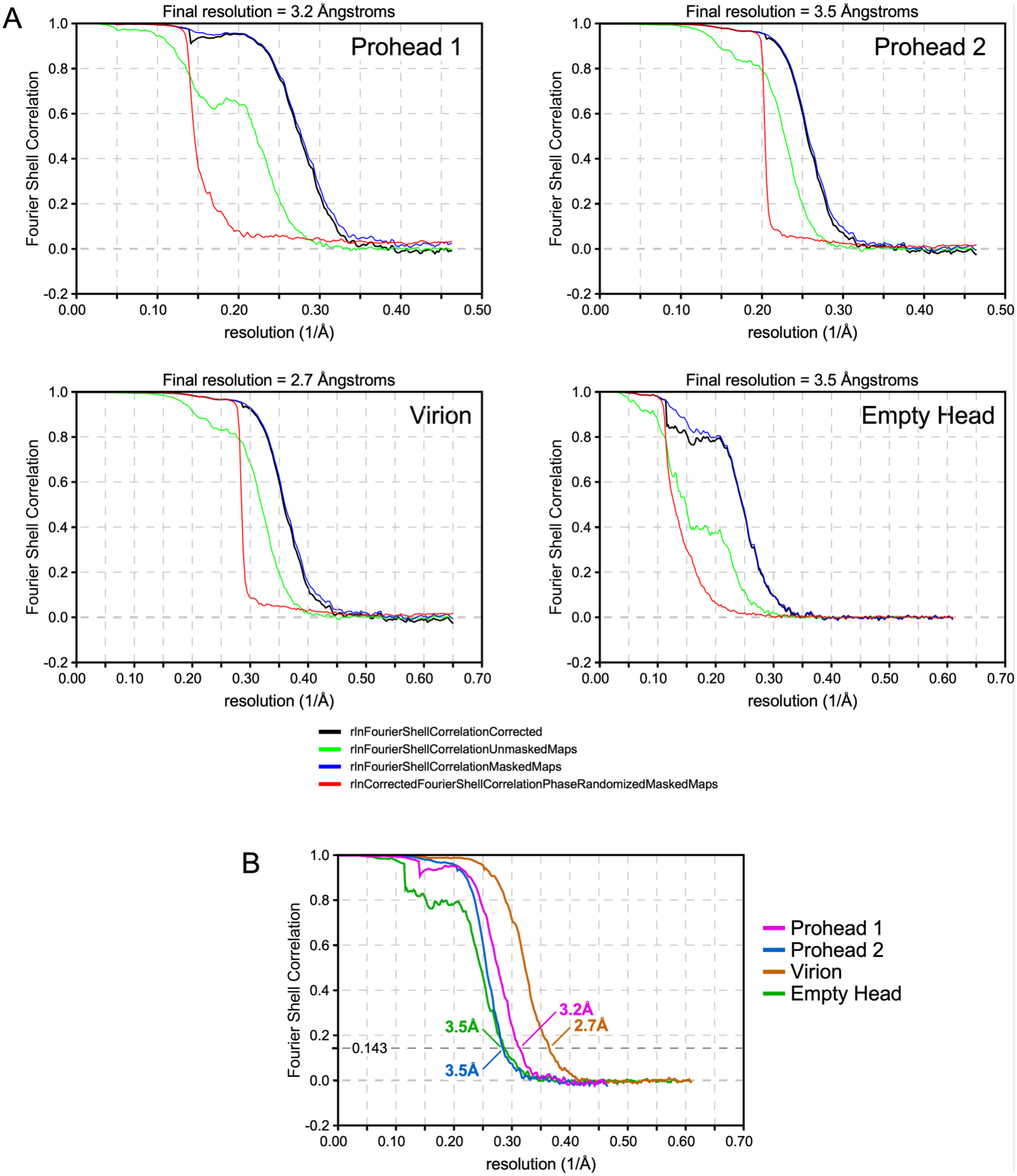


**Fig. S1.**

(A) Fourier Shell Correlation (FSC) curves for estimating resolution of each density map, as reported by Relion. (B) The FSC curves for all density maps, including resolution estimates from where the curves cross the so-called "gold-standard" correlation coefficient limit of 0.143.

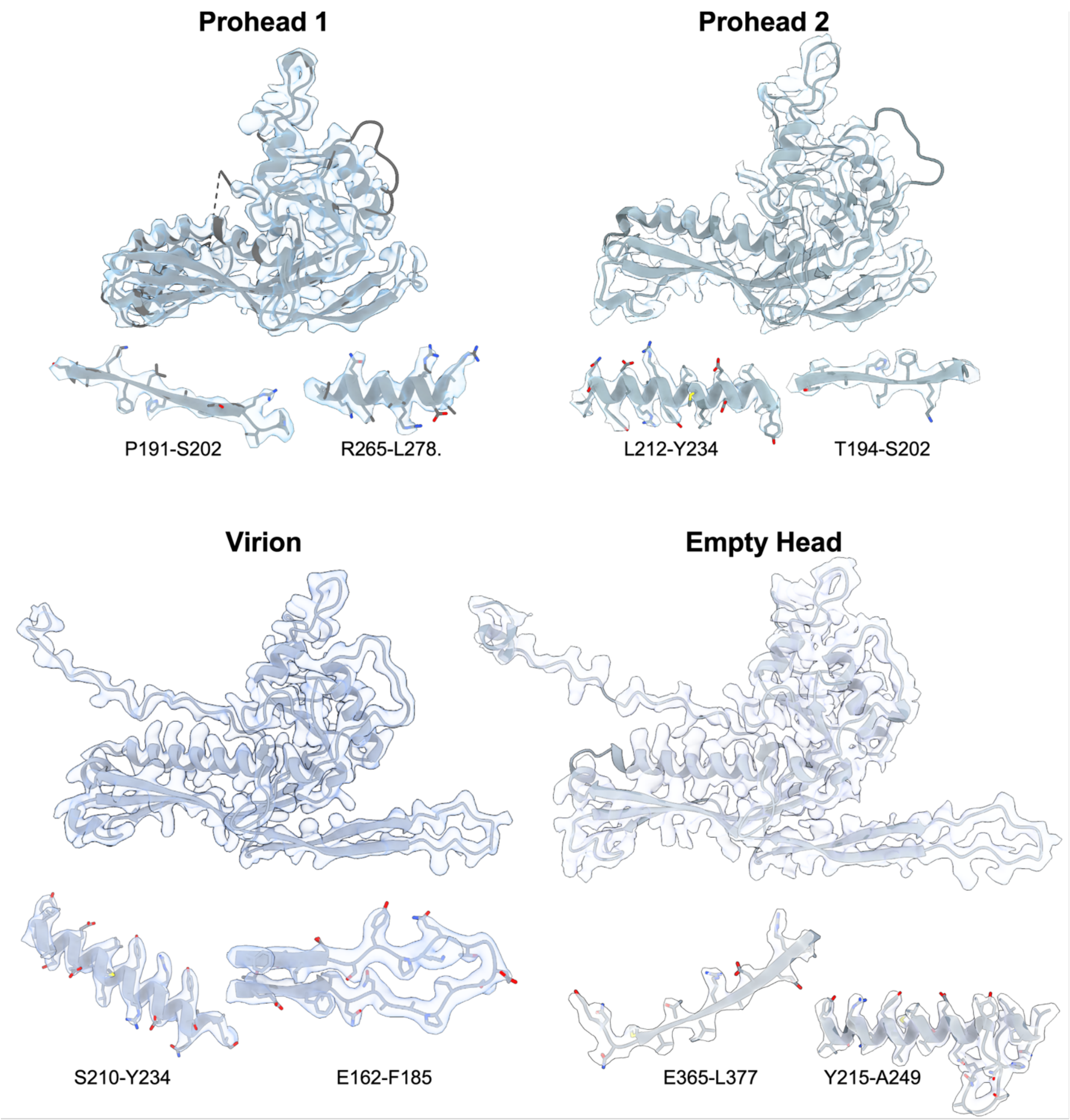


**Fig. S2.**
Atomic models of the D3 MCP derived from the four capsid forms, as indicated, including representative regions of density with sidechain fits.

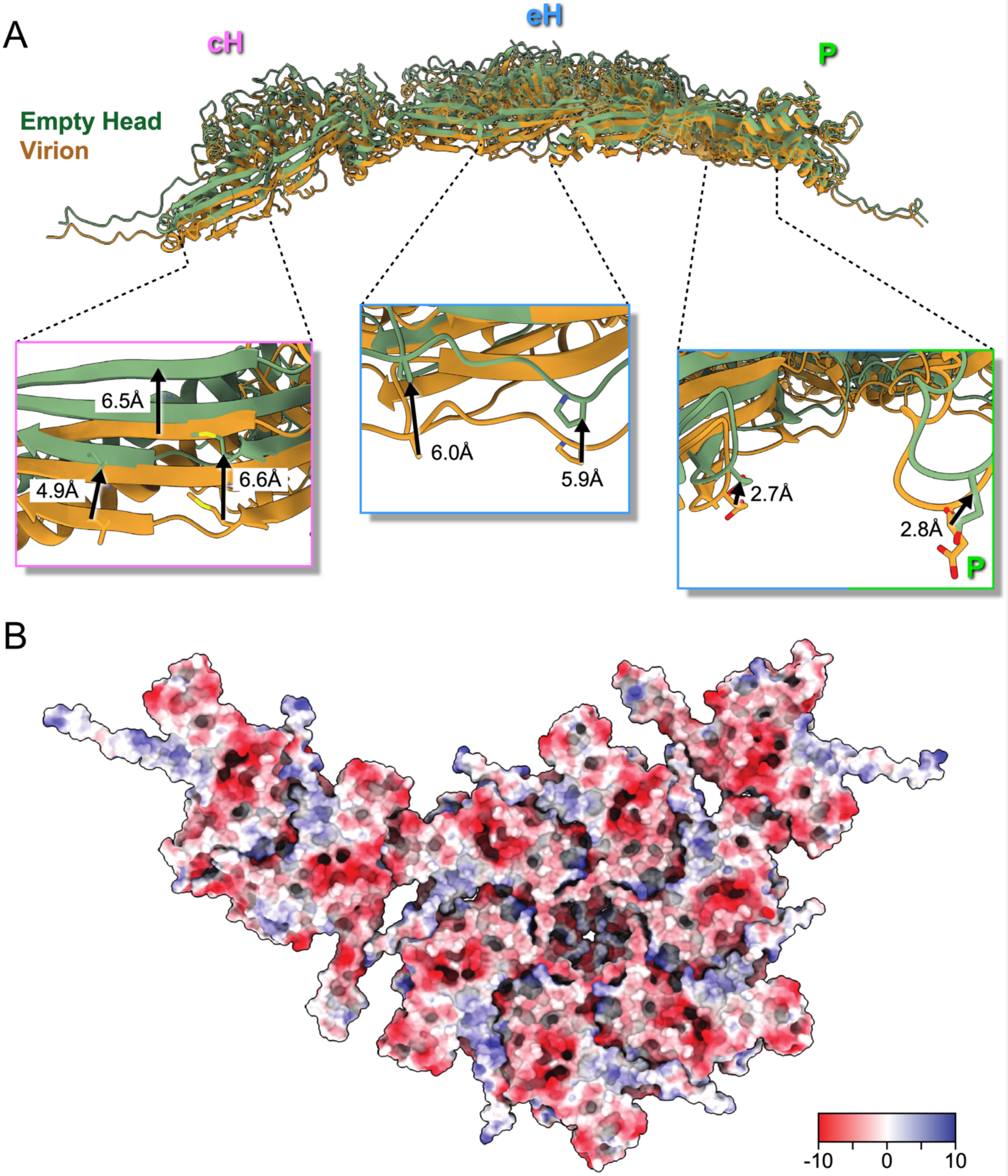


**Fig. S3.**

(A) ASU of the Virion Head and Empty Head, post-DNA ejection, demonstrating the range (~2-7Å) of radius difference between the capsid with and without packaged DNA. (B) Electrostatic potential of the virion ASU viewed from the capsid interior.

A

B

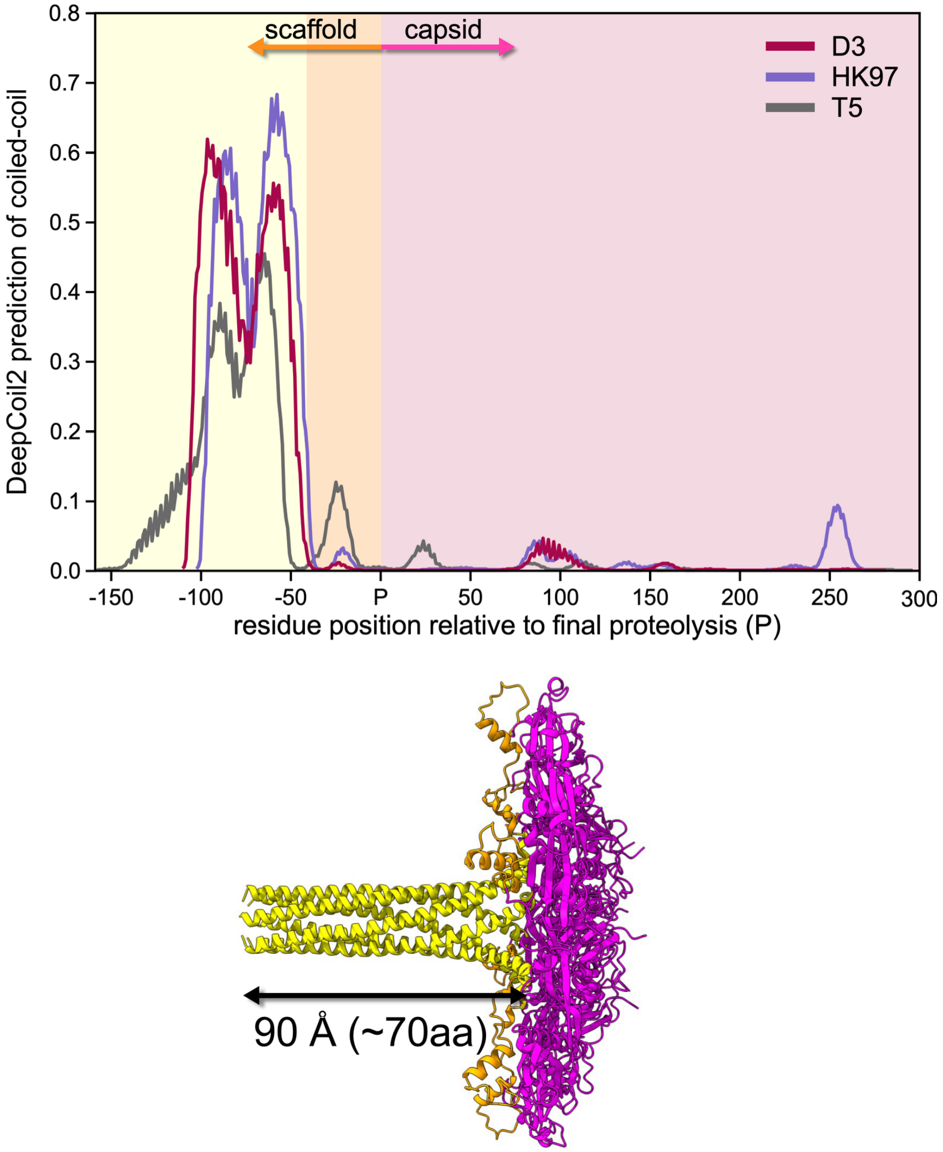


**Fig. S4.**

(A) Propensity for the MCPs of three HK97-like phages to adopt coiled-coil fold, as predicted from the amino acid sequences by DeepCoil2 (*63*). The N-terminal regions of the scaffold domains are strongly indicated to form coiled-coils, and the lengths correlate with capsid size: HK97 – scaffold domain of 101 residues, T=7; D3 – 111 residues, T=9; and T5 – 160 residues, T=13. (B) A capsomer model of five D3 MCP chains as predicted by Alphafold 2 (*64*). This predicted structure shows an extended coiled-coil fold (yellow) for the N-terminal region of the scaffold domain as well as a PDAR-like fold (orange) for the C-terminal part. The length of the coiled-coil fold (90Å) is compatible with the height of the towers of density that we observe in our prohead I map.

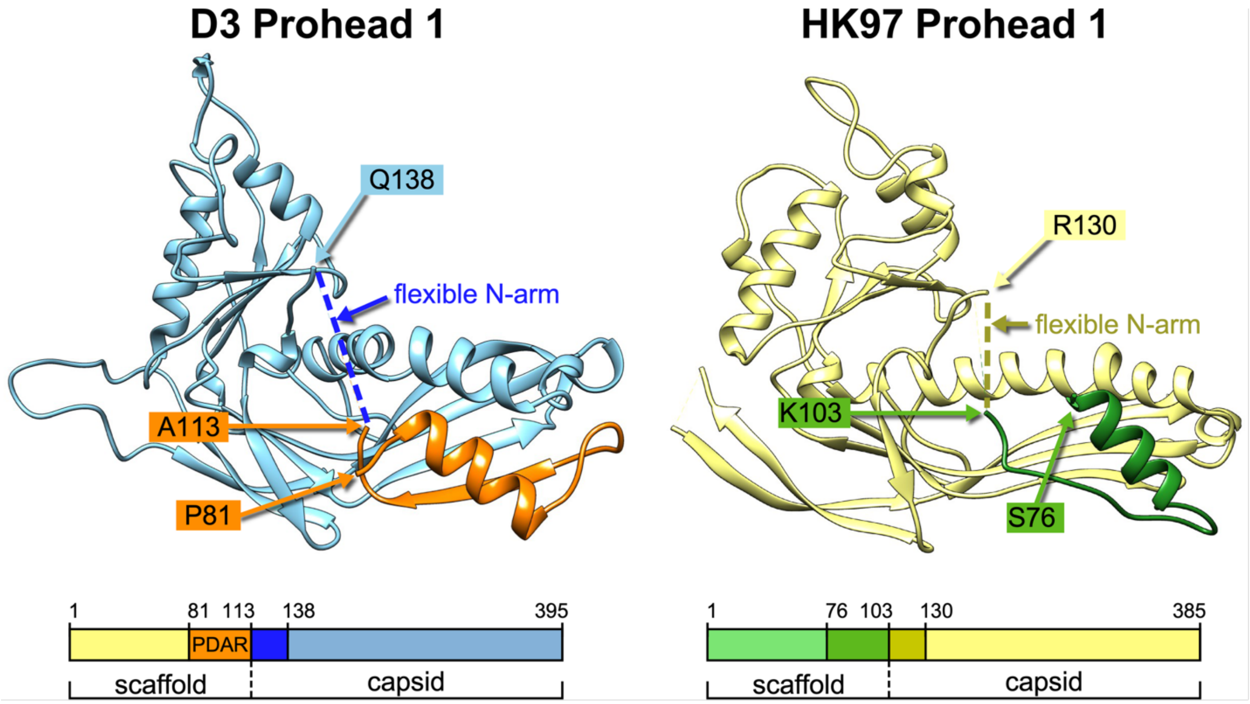


**Fig. S5.**

Comparison of the Prohead 1 subunit models of phages D3 and HK97 indicating the boundaries of regions that could be modeled confidently from cryoEM reconstructions. In both cases, the N-arm of the capsid domains was unresolved, but the C-terminal portion of the scaffold was well resolved, including the PDAR (D3: orange, HK97: dark green). Data for HK97 are from (*65*).

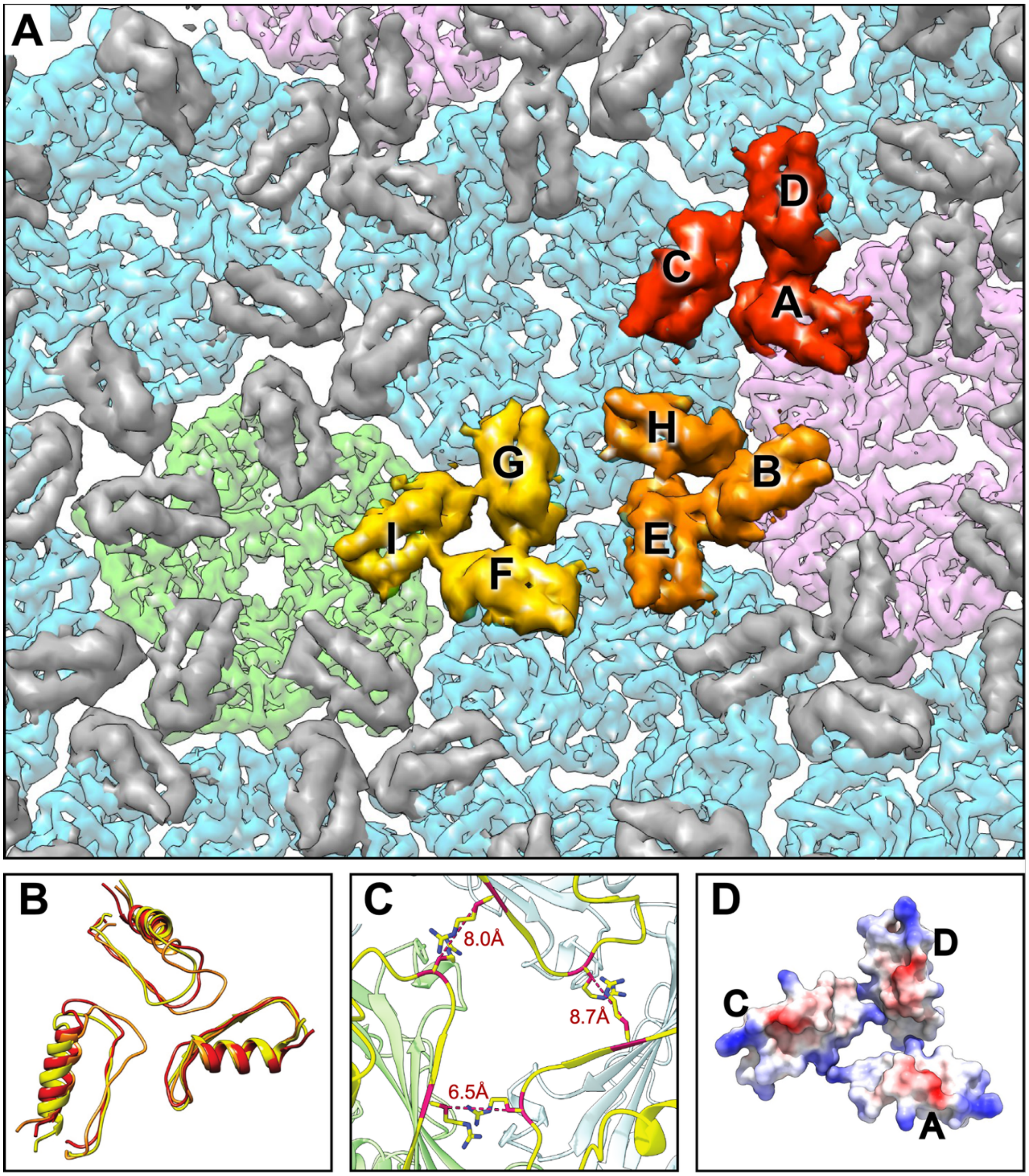


**Fig. S6.**

(A) Low resolution density map of Prohead 1 highlighting the three unique PDAR trimers at each pseudo-3-fold location (B) The three PDAR timers overlaid, exhibiting orientation differences. (C) Distance measure between the β-carbon of the interacting arginine residues on participating subunits (I,G,F) of the PDAR trimer at the penton-edge hexon interface. (D) Electrostatic rendition of the PDAR surface demonstrating an absence of attractive interactions between the three subunits.

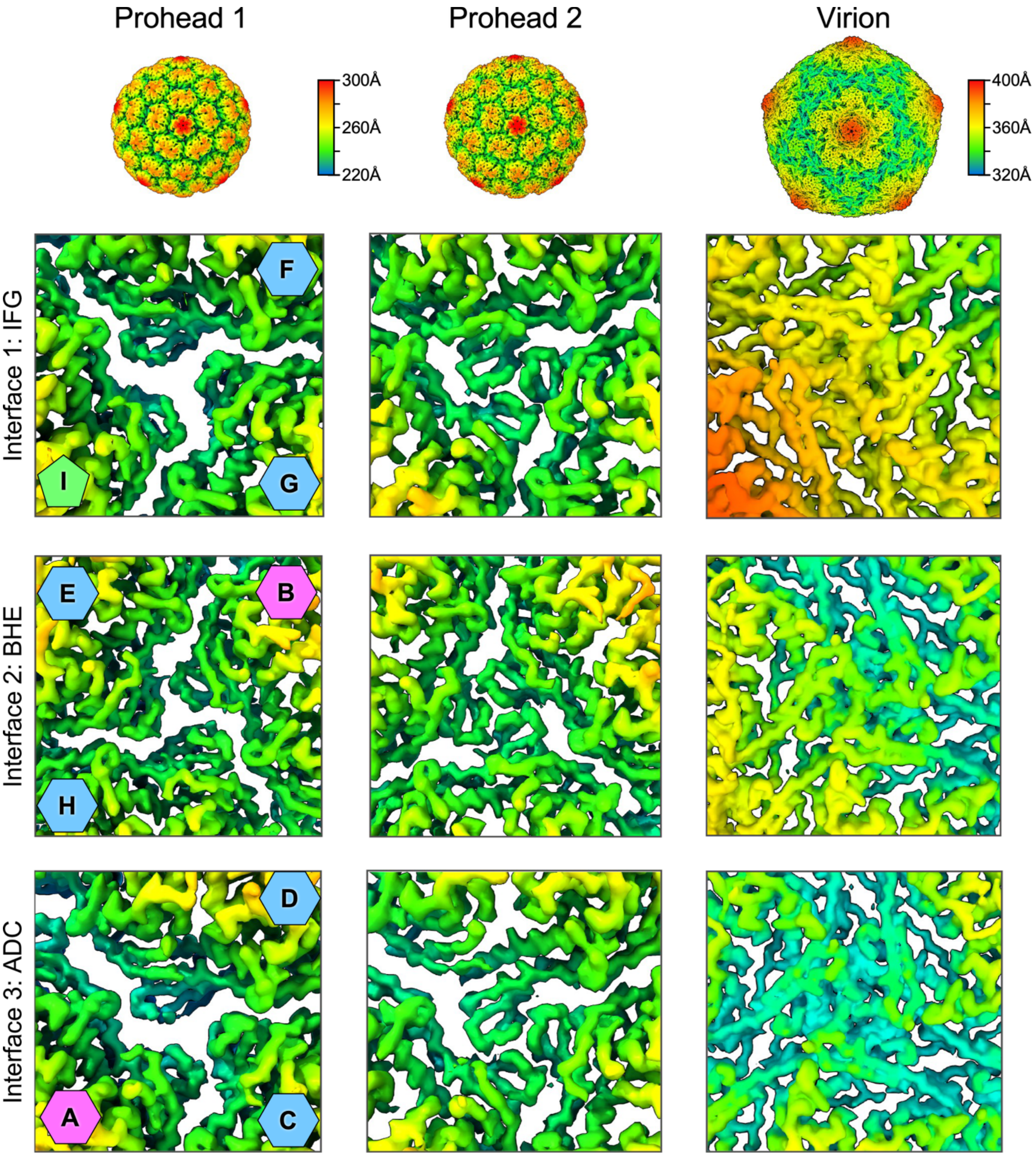


**Fig. S7.**

Comparison of density distributions at the three local 3-fold sites, as indicated at left. Surfaces are colored by radius, as shown by the scales. Note that none of the local 3-fold sites exhibits strict 3-fold icosahedral symmetry, thus subunit interactions are not identical within or between sites. Subunits are indicated by letter, and the source capsomer by the colored penton (green) and hexons (eH: blue, cH: pink), as shown in Figure 1C.

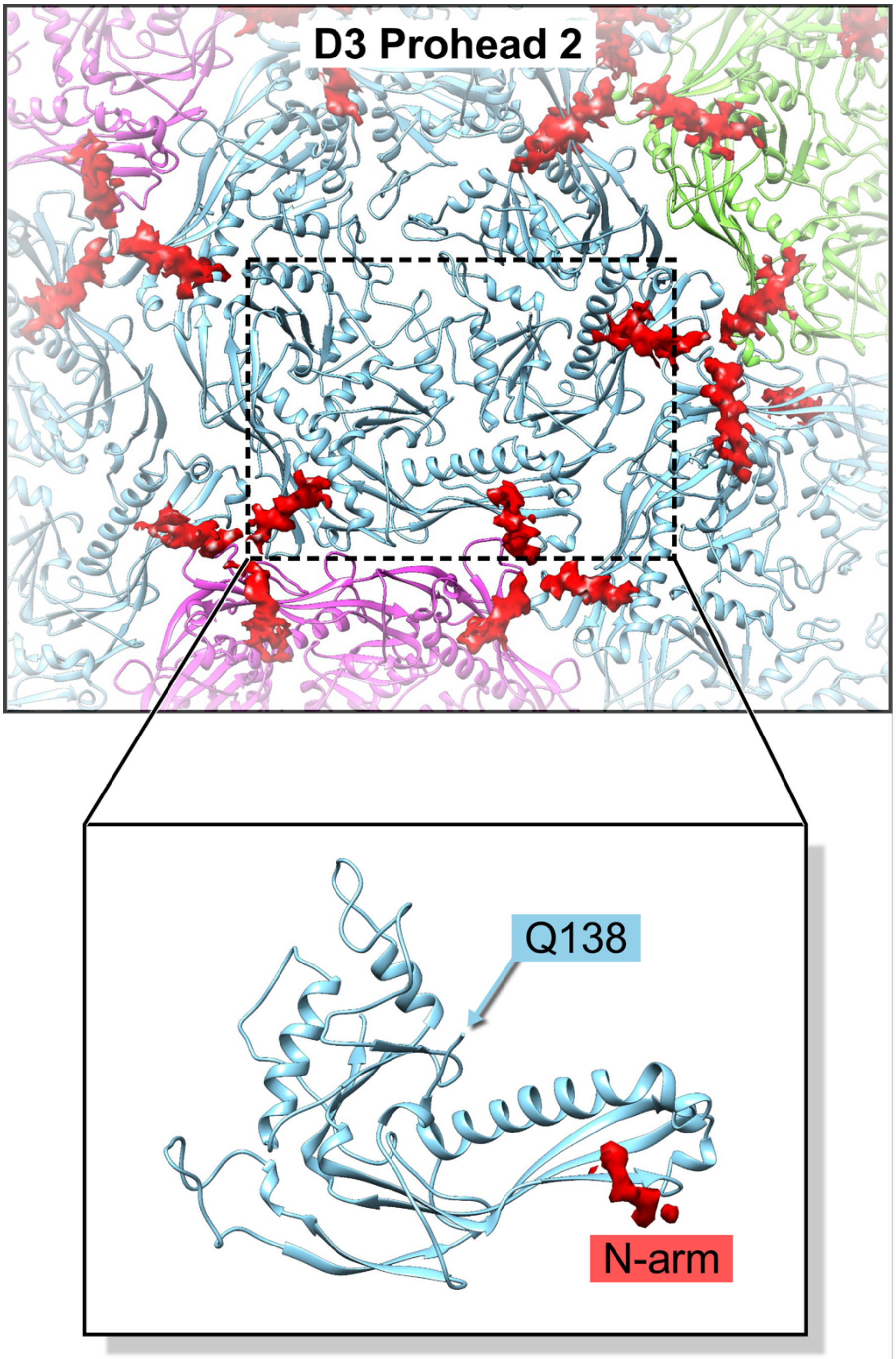


**Fig. S8.**

Unmodeled density (red) in the D3 Prohead 2 map is consistent with the location of the amino-terminal part of the N-arm based on its location near the point of scaffold cleavage.

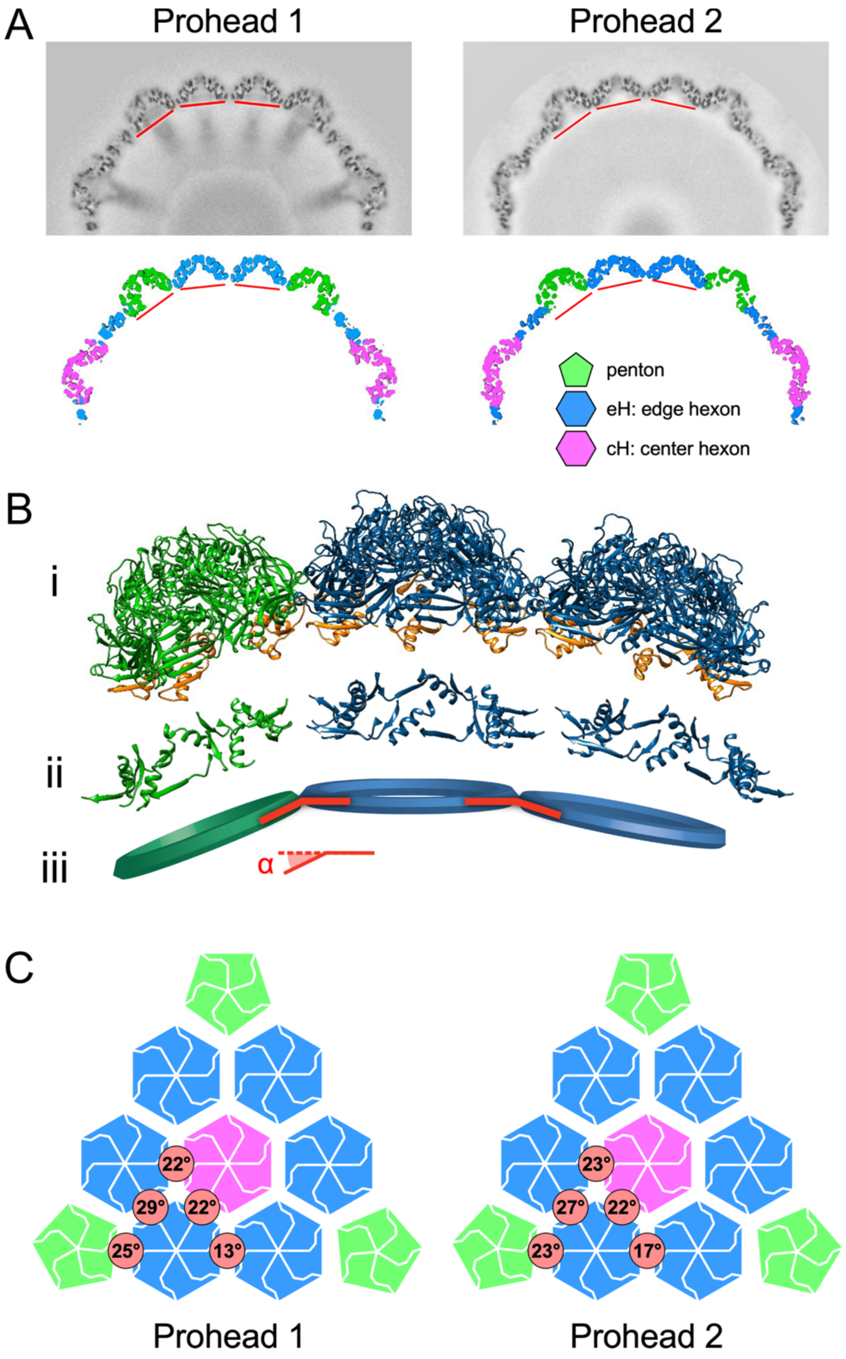


**Fig. S9.**

(A) Central sections through the Prohead 1 and Prohead 2 density maps in grey scale (top) and color-coded according to capsomer, as indicated (bottom). The red lines indicate planes through each capsomer that are defined by residues 148-154, 207-222, and 348-358, as shown in the model view (B) for Prohead 1: (i) capsomers, including PDAR domains in orange; (ii) residues defining the capsomer planes; and (iii) a schematic of how dihedral angles between capsomers are measured. Larger angles indicate greater curvature. (C) Dihedral angles measured between capsomers, as given in the red circles.

| Primer Name | Sequence |
|---|---|
| D3right1 | CGCAAGCTTGCCGGTCTCCCGGCCCCT |
| D3left1 | CGGCTCGAGCTCATGAACTGGTTCAGCCAGCCA |
| D3seq1 | GCCGTCGACAAAGGGTGGGCA |
| D3seq2 | CAGGTAAACACCCAAATCGCC |
| D3seq3 | AGTCGCTCATTGCGATTCTCC |
| d3p1 | AATGGCCAGGCCTTCGAATACGTCGC |
| d3p2 | GGCTCGGCTGGGCGCTCATTTC |
| d3p2a | AGTAGGATCCGGCTCGGCTGGGCGCTCATTTC |
| D3p-33 | CGGCTCGAGCCACCCCCGCCAGCAGCGCCA |
| D3portN | GAGTGTACGAGCGGAAAGC |
| D3portC+SphI | TAGTACGCATGCCGCTGCCGGAAGATTT |
| DP135 | GATCATCAGGAAGGCGGCGCGGC |
| DP136Q | CAGAACGCCTGGCTGATCGCCAT |
| T7phi10 | TCACCGTCATCACCGAAACGC |
| RD96-55 | TCGCTGAGATAGGTGCCTC |
| D3seq1 | GCCGTCGACAAAGGGTGGGCA |
| D3seq3 | AGTCGCTCATTGCGATTCTCC |

**Table S1.**

Primers used for this study. Sequences are given in 5' to 3' direction. pT7-5 sequence in black; D3 sequence in blue; Mutant sequence in red. T7phi10, RD96-55, d3seq1, and d3seq3 were sequencing primers.

| Name | Description | Notes | Reference |
|---|---|---|---|
| pLysS | | *p15Aori, T7 lysozyme, [$Cm^R$].* | Studier 1991 (*66*) |
| pT7-5 | | *T7 promoter[$Amp^R$].* | Vos 1988 (*67*) |
| pD3 | pT7-5::D3gene5gene6 | Primers *D3right1* and *D3left1* used in amplification of phage D3 genome insert. Digested with *XhoI* and *HindIII* and subcloned into pT7-5 vector, which was digested with SalI and HindIII. | This study |
| pD3Δ1 | pT7-5::D3gene6 | *ApoI* and *EcoRI* digest of *pD3* to delete protease gene. Ligated *ApoI->EcoRI* piece to itself. | This study |
| pD3i | pT7-5::D3gene5H141Qgene6 | *DP135* and *DP136Q* primers used for site mutagenesis with *D3right1(rev)* and *D3left1* in amplification of *pD3* insert with H141Q mutation of protease gene. Digested with *XbaI* and *SphI* and subcloned into *pD3* vector. | This study |
| pD3P | pT7-5::D3gene4gene5gene6 | Primers *d3p1* and *d3p2a* used with phage D3 genome to amplify insert. Digested with *BamHI* and *NsiI*, and subcloned into *pD3* vector. | This study |
| pD3Δ2P | pT7-5::gene4gene6 | *pD3P* digested with *EcoRI*. Ends filled with Klenow polymerase (Boehringer Mannheim, Ingelheim, Germany) and dNTPs and religated. Insert was amplified with primers *D3portN* and *D3portC+SphI* from *pD3P* Klenow filled plasmid (*pD3PK*). Digest insert with *ApoI* and *BamHI* and subclone into *pD3PK* vector. | This study |
| pD3iP | pT7-5::gene4gene5H141Qgene6 | *DP135* and *DP136Q* primers used for site mutagenesis with *D3right1(rev)* and *D3left1*in amplification of *pD3* insert with H141Q mutation of protease gene. Digested with *XbaI* and *SphI* and subcloned into *pD3P* vector. | This study |

**Table S2.**

Plasmids used for this study.

| Plasmid Name | Description | Reference |
|---|---|---|
| **For Plasmid Purification** | | |
| | DH10B | Durfee (2008) (*68*) |
| **For Protein Expression** | | |
| | BL21(DE3) | Studier 1986 (*69*) |
| pD3 | BL21(DE3) pLysS pD3 | This study |
| pD3Δ1 | BL21(DE3) pLysS pD3Δ1 | This study |
| pD3i | BL21(DE3) pLysS pD3i | This study |
| pD3P | BL21(DE3) pLysS pD3P | This study |
| pD3Δ2P | BL21(DE3) pLysS pD3Δ2P | This study |
| pD3iP | BL21(DE3) pLysS pD3iP | This study |

**Table S3.**
Bacterial strains used for in this study.

**D3 Portal Gene**

GTGAGTAAGAGTCTCGGAAAAGTCCTGAGCAGTGCTACGTCTGCGCCCAGGTCTTCATTGTTCGGCTGGGGGGTAAGACCATC
CGCCTGACAGATGGCGCGTTCTGGTCGCAGTTCCTGGGGCGAGAGTCGTCTAGCGGAAAAAAAGTCACTGTCGACAAGGCAATG
AAGCTGTCTGCGGTATGGGCTTGTGTTCGCTTGATCTCTACTTCTGTCGCCGGTCTTCCGCTTGGAGTGTACGAGCGGAAAGCG
GACGGAAGCAGAGTCGATGCTCGGTCGTTCCCGCTCTACGATGTTGTTCACAACAGCCCCAACGACGACATGACGGCCTTCCAG
TTCTGGCAGGCCATGGTCGCATCGATGCTGCTTTGGGGTAACGCATACGCGGAGATTCGCCGCGCTGCGGGCAGACCGGCTGCG
TTGGACTTCCTGCTTCCATCGAGGGTCGACCTGGAGTGTGATGAAAACGGTCGGCTGAAGTACTTCTATACGACAAAGAAGGGT
GCTCGTAGAGAGATCGAGCGTACTAACATGCTGCACATCCCGGCGTTCACGCTGGATGGTCGAATTGGTCTCTCTGCAATCAGA
TACGGCGTTGATGTCTTCGGCTCGGTCATGTCGGCGGAGGACGCAGCCAACGGCACATTCAAGAACGGACTTCTACCCACGGTC
GCATTCAAGGTTGATCGCATTCTCCAGCCTGCGCAGCGGGAGGAGTTCAGGGAGTATGTGAAGTCCGTATCGGGCGCGATGAAC
TCCGGAAGATCCCCGGTTCTGGAGCAGGGGATTACCCCTGAAACCATCGGCATCAATCCGGTCGATGCTCAGTTGCTGGAGACG
CGAGAGCATGGCGTGATCGAGATTTGCAGATGGTTCGGGGTGCCGCCCTGGATGATCGGCCAGACCGACAAGGGGAGCAACTGG
GGGACAGGGCTTGAACAGCAGATGCTCGCGTTCCTGACATTCTCGATCAGCTCGATCACCAATCAGATTCAGCAGTGCGTCAAC
AAGCGGCTGCTAACTGCGCCCGAGCGGATTCGCTATTACGCCGAGTTCTCACTTGAGGGGTTCCTGAAGGCTGATAGCGCTGGT
CGCGCTGCCTGGTACAGCACTATGGCGCAAAACGGTTTCATGACCCGCAACGAAGGTCGCCGGAAAGAGAACCTGCCAGAACTC
CCCGGCGGAGACATTCTCACCGTCCAATCCAACCTAGTTCCAATCGACCAACTCGGTCAATCTAACGAGAGCCAGGCCGTCCGC
GCCGCGCTCATGAACTGGCTCAGCCAGCCAGAACCACAGGAGTAA

**D3 Protease Gene**

ATGACTCTGCGAAATCTTCCGGCAGCGCCGGAGGCTCGCCCGCGCTCGGGCGTCCAGTGCGACCTGGCGCCCAAAGCGCTAGAT
GCATGGCGTCCTGAGCTTCGAGCAGCTTCTGGCGATAACCCGGACTCCACGATCACCATCTACGAGCCGATTGGCTACGACTGG
TGGACCGGTGAAGGTGTCACGGCAAAACGCATTGCTGGCGCTCTGCGCTCCATCGGCAACGATGTCGATGTGACCGTGAACATC
AACAGCCCCGGCGGCGACGTATTCGAAGGCCTGGCCATTTACAACCTGCTGCGCGAGCACAAGGGCAAGGTCACGGTGAACATC
ATCGGCCTGGCTGCCTCTGCCGCCTCTTTCATCGCCATGGCGGGGGATGAAATCCGCATCGGCCGCGCCGCCTTCCTGATGATC
CATAACGCCTGGCTGATCGCCATGGGTAATCGGAATGATCTCCGTGAGATAGCCGATTGGCTGGAGCCATTCGACATGACGCTG
GCTGACATTTACGCACAGCGCACGGAAATCGACATCGACGACATCGTGAAACAGATGGACGCCGAGACCTGGATCGGCGGGCGC
GAAGCCGTCGACAAAGGGTGGGCAGATGCCTTCCTGGAGTCCGACGAGATCTCCAGCGCTCCCAGCAACCGCAGCGAAGCCATC
CTGGCCAAGCGCCGGATGGATGCCGCCCTGGCTCGCAGCGGAATGCCGCGAAGCCAGCGCAATGAACTCATCAACGACTTCAAG
ACCAGCATGCTTGGCGCTGCTGGCGGGGGTGGTGACACCCCGACCGATATGCCTGGCGCTGTCGCTCCTGACCTCTCCGCTGCA
CTACGGGCAGCACAAGACATCACCAAATTCCTCCAAGGAGAATCGCAATGA

**D3 Major Capsid Gene**

ATGAGCGACTTCGAAAAACAAATCGGCGAACTGAACGCCAGCCTCAAGCAGGTCGGGGACCAAATCAAGTCCCAGGCCGAACAG
GTAAACACCCAAATCGCCAATTTCGGCGAAATGAACAAGGAAACCCGCGCCAAGGTCGACGAACTGCTGACTGCCCAGGGCGAA
CTGCAAGCACGACTGAGCGCCGCGGAACAAGCCATGCTGGCCAACGAGAAGCGTGACGGCGGCGAGGAAGCACCGAAGACCGCC
GGCCAAATGGTCGCAGAGAGCCTGAAAGAGCAGGGTGTAACCAGCTCCCTGCGCGGCTCGCATCGCGTATCCATGCCGCGCTCG
GCCATCACCTCCATCGACGGCTCTGGCGGCGCCCTGGTGGCTCCTGATCGTCGCCCCGGTGTCGTTGCCGCTCCGCAGCGTCGA
CTGACCATCCGCGACCTGGTTGCGCCTGGCACCACTGAGTCGAACTCCGTCGAGTACGTCCGCGAGACCGGCTTCGTCAACAAT
GCCGCTCCTGTTTCGGAAGGCACCCAGAAGCCATACTCCGACCTGACCTTCGAACTGGAAAACGCGCCGGTTCGCACCATCGCA
CACCTGTTCAAGGCAAGTCGCCAGATCCTGGACGACGCTTCGGCCTTGCAGAGCTACATCGATGCGCGCGCTCGTTACGGCCTG
ATGCTGGTCGAAGAAGGTCAACTGCTCTACGGGAACGGGACCGGCGCCAATCTGCACGGCATCATTCCGCAGGCACAGGCCTAC
GCGCCGCCGAGTGGCGTAGTGGTAACCGCCGAGCAGCGAATCGACCGCATCCGCCTGGCGATCCTTCAGGCGCAACTGGCCGAG
TTCCCGGCCAGCGGTATCGTGCTCAACCCCATCGACTGGGCGCTGATCGAGCTGACCAAGGACGCCGAGAACCGTTACATCATC
GGCAGCCCGCAGAACGGCACCACTCCGACCCTCTGGCGTCTGCCGGTGGTGGAAACCCAGGCCATCACTCAGGACGAGTTCCTG
ACCGGTGCGTTCTCTCTCGGCGCCCAGATCTTCGACCGCATGGACATCGAGGTTCTGGTTTCCACCGAGAACGACAAGGACTTC
GAGAACAACATGGTCACCATCCGCGCTGAGGAGCGGCTGGCCTTCGCGGTCTATCGCCCCGAGGCTTTCGTGACTGGTTCGCTG
ACCGCCAGCTAA

**Table S4.**

Gene Sequences. Notes:

- Sanger sequencing of plasmids containing D3 genes showed discrepancies compared with reported sequence of the D3 genome (Genbank: AF165214) by Kropinski (*70*). Shown is corrected sequence per Sanger sequencing results with discrepancies highlighted in red.
- 6 errors in the published D3 sequence have been confirmed and led to nucleotide changes in the coding sequences. They are (erroneous nucleotide in red):
  - AAG->GAG (3506) and TTC->CTC (3542) in gene 4
  - ATT->ACT (3576) and GTT->GCT (3606) in gene 5
  - TGT->GGT (5106) and AAC->ACC (5354) in gene 6.
- Our revised genome sequence was deposited to Genbank (PV029250).

| |
|---|
| **D3 Portal Protein**<br>MSKSLGKVLSSATSAPRSSLFGWGGKTIRLTDGAFWSQFLGRESSSGKKVTVDKAMKLSAVWACVRLISTSVAGLP<br>LGVYERKADGSRVDARSFPLYDVVHNSPNDDMTAFQFWQAMVASMLLWGNAYAEIRRAAGRPAALDFLLPSRVDLE<br>CDENGRLKYFYTTKKGARREIERTNMLHIPAFTLDGRIGLSAIRYGVDVFGSVMSAEDAANGTFKNGLLPTVAFKV<br>DRILQPAQREEFREYVKSVSGAMNSGRSPVLEQGITPETIGINPVDAQLLETREHGVIEICRWFGVPPWMIGQTDK<br>GSNWGTGLEQQMLAFLTFSISSITNQIQQCVNKRLLTAPERIRYYAEFSLEGFLKADSAGRAAWYSTMAQNGFMTR<br>NEGRRKENLPELPGGDILTVQSNLVPIDQLGQSNESQAVRAALMNWLSQPEPQE |
| **D3 Protease Protein**<br>MTLRNLPAAPEARPRSGVQCDLAPKALDAWRPELRAASGDNPDSTITIYEPIGYDWWTGEGVTAKRIAGALRSIGN<br>DVDVTVNINSPGGDVFEGLAIYNLLREHKGKVTVNIIGLAASAASFIAMAGDEIRIGRAAFLMIHNAWLIAMGNRN<br>DLREIADWLEPFDMTLADIYAQRTEIDIDDIVKQMDAETWIGGREAVDKGWADAFLESDEISSAPSNRSEAILAKR<br>RMDAALARSGMPRSQRNELINDFKTSMLGAAGGGGDTPTDMPGAVAPDLSAALRAAQDITKFLQGESQ |
| **D3 Major Capsid Protein**<br>MSDFEKQIGELNASLKQVGDQIKSQAEQVNTQIANFGEMNKETRAKVDELLTAQGELQARLSAAEQAMLANEKRDG<br>GEEAPKTAGQMVAESLKEQGVTSSLRGSHRVSMPRSAITSIDGSGGALVAPDRRPGVVAAPQRRLTIRDLVAPGTT<br>ESNSVEYVRETGFVNNAAPVSEGTQKPYSDLTFELENAPVRTIAHLFKASRQILDDASALQSYIDARARYGLMLVE<br>EGQLLYGNGTGANLHGIIPQAQAYAPPSGVVVTAEQRIDRIRLAILQAQLAEFPASGIVLNPIDWALIELTKDAEN<br>RYIIGSPQNGTTPTLWRLPVVETQAITQDEFLTGAFSLGAQIFDRMDIEVLVSTENDKDFENNMVTIRAEERLAFA<br>VYRPEAFVTGSLTAS |

**Table S5.**

Protein Sequences. Notes:

- 6 errors in the published D3 sequence have been confirmed and led to amino acid changes in the coding sequences. They are (erroneous nucleotide in red):
  - K415E and F427L in gene 4
  - I2T and V12A in gene 5
  - C230G and N299T in gene 6

**Movie S1.**

From Prohead 1 to Prohead 2. Comparison of surface of Prohead 1 to Prohead 2 showing that after removal of the scaffold the capsomers are subject to reshaping, notably at the 3fold interface.

**Movie S2.**

Structural transition from the Prohead 1 to Prohead 2 at the pseudo-3-fold (IFG interface). Related to Fig. 7, the movie illustrates reorganization of the interface involving residue F364. The residue is proximal to the scaffold residue R102 in Prohead 1 and, following scaffold removal by proteolysis, F364 is displaced toward the center of the pseudo-3-fold, sandwiched by residues N360 and R372 in Prohead 2.

**Movie S3.**

Structural transition from Prohead 2 to the expanded head at the pseudo-3-fold (IFG interface). Related to Fig. 8, the movie illustrates the stability of the N360-F364-R372 cluster during capsid expansion.

**Movie S4.**

An animated view of 72 D3 subunits expanding shows the preservation of a salt bridge between E162 on the base of each E-loop and R219 on the backbone helix of the adjacent subunit within the same capsomer. The movie features linear morphing between models of Prohead 1, Prohead 2, and Head with pauses after each transition. Subunits are shown as ribbons; the nine D3 asymmetric unit subunits are in bold colors. E-loops are green, G-loops orange, P-loops magenta; E162 (red) and R219 (aqua) atoms are shown as space-filled spheres. Captions on the right indicate assembly stages. Prohead 1 and 2 models include placeholder residues at the E-loop tips (due to insufficient density) to facilitate morphing, though these are absent from the deposited models. The D3 MCP N-arms were not modeled in Proheads 1 or 2 and appear abruptly mid-movie in the mature capsid. Morphing was done with VMD 1.92 (*71*), saved as DCD-format trajectories, opened in Chimera (*72*) via MD Movie, and exported as .png frames using Movie Recorder. Final editing, including captions, was done in Blender Video Editor (version 3.1.2, http://www.blender.org) and rendered to mp4.
